# SBMLKinetics: A Tool for Annotation-Independent Classification of Reaction Kinetics for SBML Models

**DOI:** 10.1101/2022.11.17.516936

**Authors:** Jin Xu

## Abstract

**Background:** Reaction networks are widely used as mechanistic models in systems biology to reveal principles of biological systems. Reactions are governed by kinetic laws that describe reaction rates. Selecting the appropriate kinetic laws is difficult for many modelers. There exist tools that attempt to find the correct kinetic laws based on annotations. Here, I developed annotation-independent technologies that assist modelers by focusing on finding kinetic laws commonly used for similar reactions.

**Results:** Recommending kinetic laws and other analyses of reaction networks can be viewed as a classification problem. Existing approaches to determining similar reactions rely heavily on having good annotations, a condition that is often unsatisfied in model repositories such as BioModels. I developed an annotation-independent approach to find similar reactions via reaction classifications. I proposed a two-dimensional kinetics classification scheme (2DK) that analyzed reactions along the dimensions of kinetics type (K type) and reaction type (R type). I identified approximately ten mutually exclusive K types, including zeroth order, mass action, Michaelis-Menten, Hill kinetics, and others. R types were organized by the number of distinct reactants and the number of distinct products in reactions. I constructed a tool, SBMLKinetics, that inputted a collection of SBML models and then calculated reaction classifications as the probability of each 2DK class. The effectiveness of 2DK was evaluated on BioModels, and the scheme classified over 95% of the reactions.

**Conclusions:** 2DK had many applications. It provided a *data-driven* annotation-independent approach to recommending kinetic laws by using type common for the kind of models in combination with the R type of the reactions. Alternatively, 2DK could also be used to alert users that a kinetic law was unusual for the K type and R type. Last, 2DK provided a way to analyze groups of models to compare their kinetic laws. I applied 2DK to BioModels to compare the kinetics of signaling networks with the kinetics of metabolic networks and found significant differences in K type distributions.

## Background

Reaction networks are widely used in systems biology as mechanistic models that can reveal governing principles of biological systems. The choice of kinetic laws for reactions (e.g., mass action kinetics, Michaelis-Menten kinetics) is crucial for the development of mechanistic models of biological systems. Computational studies of systems biology often involve the analysis of synthetically generated networks (e.g., [1, 2]). As noted in [3, 4], the choice of kinetic laws for these networks is as important as the set of reactions (the reactants, products, and modifiers for reactions) since reactions alone are not sufficient to determine dynamics [5]. Indeed, many computational studies only consider mass action kinetics, which may be inadequate [1]. In addition, kinetics are crucial considerations in parameter inference and model identification [6].

There is a wide diversity of kinetic laws. For example, allosteric and cooperative kinetics are important considerations in some biological systems [7, 8], and there are generalizations of these laws [9, 10], such as convenience kinetics [11]. Complications arise if some inhibitors or activators target the active site (non-allosteric). A further consideration is an order in which elemental reactions occur to form complexes such as compulsory order and ping-pong [12, 13, 14].

A significant motivation for this work was addressing the challenges modelers face with choosing appropriate kinetic laws. A detailed example of choosing kinetic laws is given for the models of glycolysis [15] and Trypanosoma models [16, 17]. Further, King-Altman [18] considers a much more complex reaction mechanism than deriving Michaelis-Menten kinetics using the steady-state assumption. Kinetic laws are chosen based on the best knowledge of the underlining kinetic mechanism. Some books describe kinetic laws, such as Segel’s enzyme kinetics [12] and others [13, 14].

Despite the examples, there remains a significant challenge with applying the theory in practice, especially for less experienced modelers. To address this, programs such as COPASI [19] and CellDesigner [20], provide pre-defined lists of kinetic laws. SBMLsqueezer 2 [21] goes further by making recommendations for kinetic laws in two ways. One way is a classification scheme that “depends on the annotations associated with each reactant, reaction and modulation”, which are “incorporated into the Systems Biology Markup Language (SBML) [22] specifications in the form of Systems Biology Ontology (SBO) [23]” [24]. “The most important sources of information in determining these categories are Minimal Information Required In the Annotation of Models (MIRIAM) [25] and SBO annotations” [21]. The other way also depends on annotations to match reactions in the SBAIO-RK [26] database to obtain kinetic laws. I emphasize that these two ways are limited because of the heavy dependence on annotations. Further, the authors note that even with annotations, there are serious limitations in that “the extraction from SABIO-RK depends on existing biochemical data and might therefore not always yield results.” Here, I expanded one aspect of the first way taken in SBMLsqueezer 2 to find similar reactions, whose core is to classify reactions. In SBMLsqueezer 2, this relies on annotations which include SBO and MIRIAM.

In this article, I developed a classification scheme that did not depend on annotations. Existing approaches to determining similar reactions rely heavily on having good annotations of chemical species, a condition that is often unsatisfied in model repositories such as BioModels. This article introduced **2DK**, a new scheme for classifying reactions based on a two dimension structure: kinetics type (**K type**) and reaction type (**R type**). K types were mutually exclusive kinds of kinetic laws such as zeroth order, mass action, Michaelis-Menten, and Hill kinetics. R types classified reactions by the number of distinct reactants and products. I developed a Python-based cross-platform tool, SBMLKinetics, that constructed reaction classifications by inputting a collection of models and outputting the probability of each **2DK** class (combinations of **K type** and **R type**).

I validated the **2DK** classification scheme using BioModels Database. I further described how **2DK** could provide a *data-driven* approach, with BioModels Database as an example, to recommending kinetic laws and providing an alert of unusual kinetic laws. Finally, I applied BioModels Database as an example again to compare the kinetics of signaling networks with the kinetics of metabolic networks. The results showed that the signaling networks were dominated by uni-directional mass action kinetics whereas metabolic networks had a proportionally larger number of regulatory kinetics in the form of kinetic laws that had a fractional representation. This is reasonable because metabolic models are derived heavily from underlying enzyme mechanisms whereas signaling pathways are generally simpler binding/unbinding reactions or gross simplifications of the kinetics because very little is known about the mechanism.

## Methods

I developed an annotation-independent classification scheme for kinetic laws based on patterns in existing reaction networks. The focus was on the standard community of SBML. The approach classified reactions along two dimensions. The first was a characterization of the algebraic expression of the kinetic law. I referred to this as the kinetics type (**K type**) of the reaction. The second was a simplification of reaction stoichiometry, namely the number of distinct reactants and the number of distinct products. I referred to this as the reaction type (**R type**) of the reaction.

### 1. Define the two-dimensional kinetics classification scheme (**2DK**)

The two-dimensional kinetics classification scheme (**2DK**) organized reactions along two dimensions: kinetics type (**K type**) and reaction type (**R type**).

#### 1.1 Define the kinetics type (K type)

On one hand, **K type** was defined mainly by the algebraic expressions of kinetic laws. For example, the kinetics with a reaction of *A* + *B* → *C* and with its kinetic law *K*_1_ is different from the same reaction *A* + *B* → *C* but with another kinetic law *K*_1_ *· A · B*. On the other hand, **K type** also included some information from the reactions. For example, it was necessary to consider whether a species inside the kinetic law was a reactant or not. To evaluate the kinetics statistically, I classified the kinetics into commonly used and mutually exclusive ten types with defined categories as below, referring to Ontology Search (OLS) [27].

- **Zeroth order (ZERO)**: the number of species in the kinetic law was zero;
- **Uni-directional mass action (UNDR)**: the kinetic law was a single product of terms and all species in the kinetic law were reactants;
- **Uni-term with the moderator (UNMO)**: the kinetic law was a single product of terms and not UNDR, namely at least one species in the kinetic law was *not* a reactant;
- **Bi-directional mass action (BIDR)**: the kinetic law was the difference between two products of terms, and the first product of terms contained species as all the reactants while the second product of terms contained species as all the products;
- **Bi-terms with the moderator (BIMO)**: the kinetic law was the difference between two products of terms and NOT BIDR;
- **Michaelis-Menten kinetics without explicit enzyme (MM)**: the kinetic law was in the format of Michaelis-Menten expressions without an explicit enzyme;
- **Michaelis-Menten kinetics with an explicit enzyme (MMCAT)**: the kinetic law was in the format of Michaelis-Menten expressions with an explicit enzyme;
- **Hill equation (HILL)**: the kinetic law was in the format of Hill equations;
- **Fraction format other than MM, MMCAT, and HILL (FR)**: the kinetic law was in the format of fraction with at least one species in the denominator and NOT MM, MMCAT, or HILL;
- **Not classified (NA)**: not classified kinetics.

Table 1 provided an example for each certain **K type**. In particular, the type FR included generalized kinetics with the kinetic law in the format of fraction based on but beyond Michaelis-Menten and Hill kinetics. All the information in Table 1 other than **K type** was from the SBML files in the BioModels Database. There are four columns: “K Type”, “BIOMD” (BioModel identifier number), “Reaction” and “Kinetic law”. In the column of “BIOMD”, i.e., 5 means BIOMD0000000005. In the “Kinetic law” column, all the species in the kinetic law are in bold. As a comparison, all the parameters are not in bold. For instance, variable compartment sizes were considered as parameters. The K types did not consider the values of the parameters; therefore, units were not considered here either.

**Table 1.**
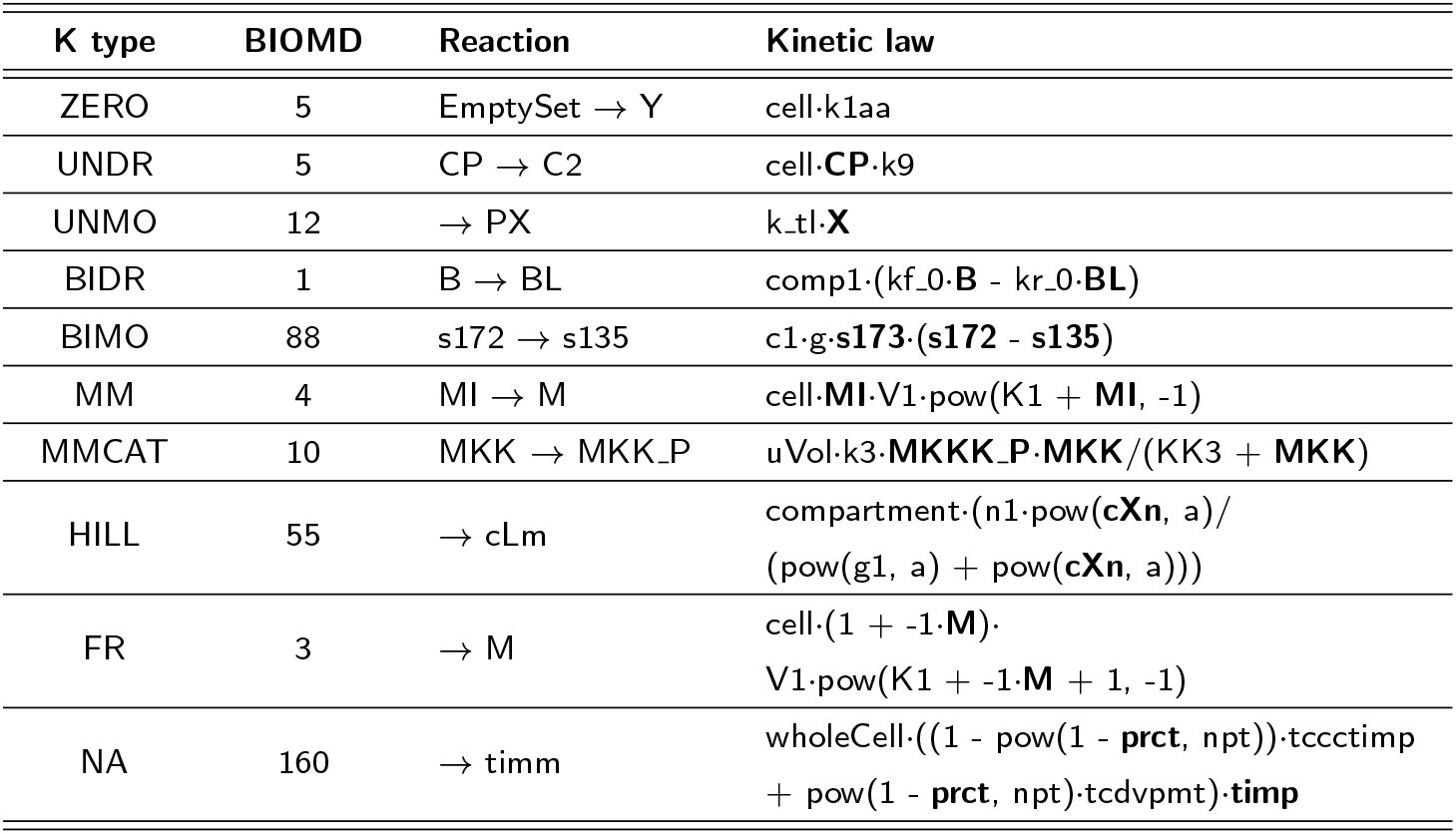
Kinetics examples for ten defined kinetics types (K type).

OLS lists the mathematical expressions of kinetic laws in SBO as four types: mass action kinetic laws, modular kinetic laws, enzymatic kinetic laws, and Hill-type kinetic laws. Table 2 described how the defined K types correspond to the mathematical expressions of kinetic laws in SBO.

**Table 2.**
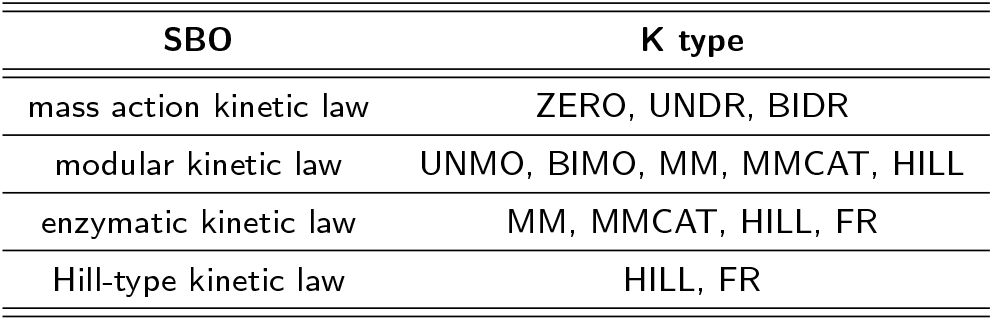
Comparison between kinetics types (K types) and Systems Biology Ontology (SBO) in Ontology Search (OLS).

| SBO | K type |
| --- | --- |
| mass action kinetic law | ZERO, UNDR, BIDR |
| modular kinetic law | UNMO, BIMO, MM, MMCAT, HILL |
| enzymatic kinetic law | MM, MMCAT, HILL, FR |
| Hill-type kinetic law | HILL, FR |

#### 1.2 Define the reaction type (R type)

**R type** was defined by the structure of reactions. It was a simplification of reaction stoichiometry in that only the number of distinct reactants and products were considered. For example, two reactants and one product are in the reaction of *A* + *B* → *C*. I defined **R type** quantitatively with a pair of the number of reactants (R = 0, 1, 2, *>*2) and products (P = 0, 1, 2, *>*2). As a comparison, the stoichiometry matrix indicates reactants and products by negative and positive signs respectively. The **2DK** classification was a combination of the **K type** and the **R type**.

### 2. Code structure and statistical analysis

I developed a Python package called SBMLKinetics that read a collection of SBML models and then did statistical analysis for the **2DK** clarifications. There were four major workflow steps in SBMLKinetics implemented by three Python scripts as shown in Figure 1.

**Figure 1.**
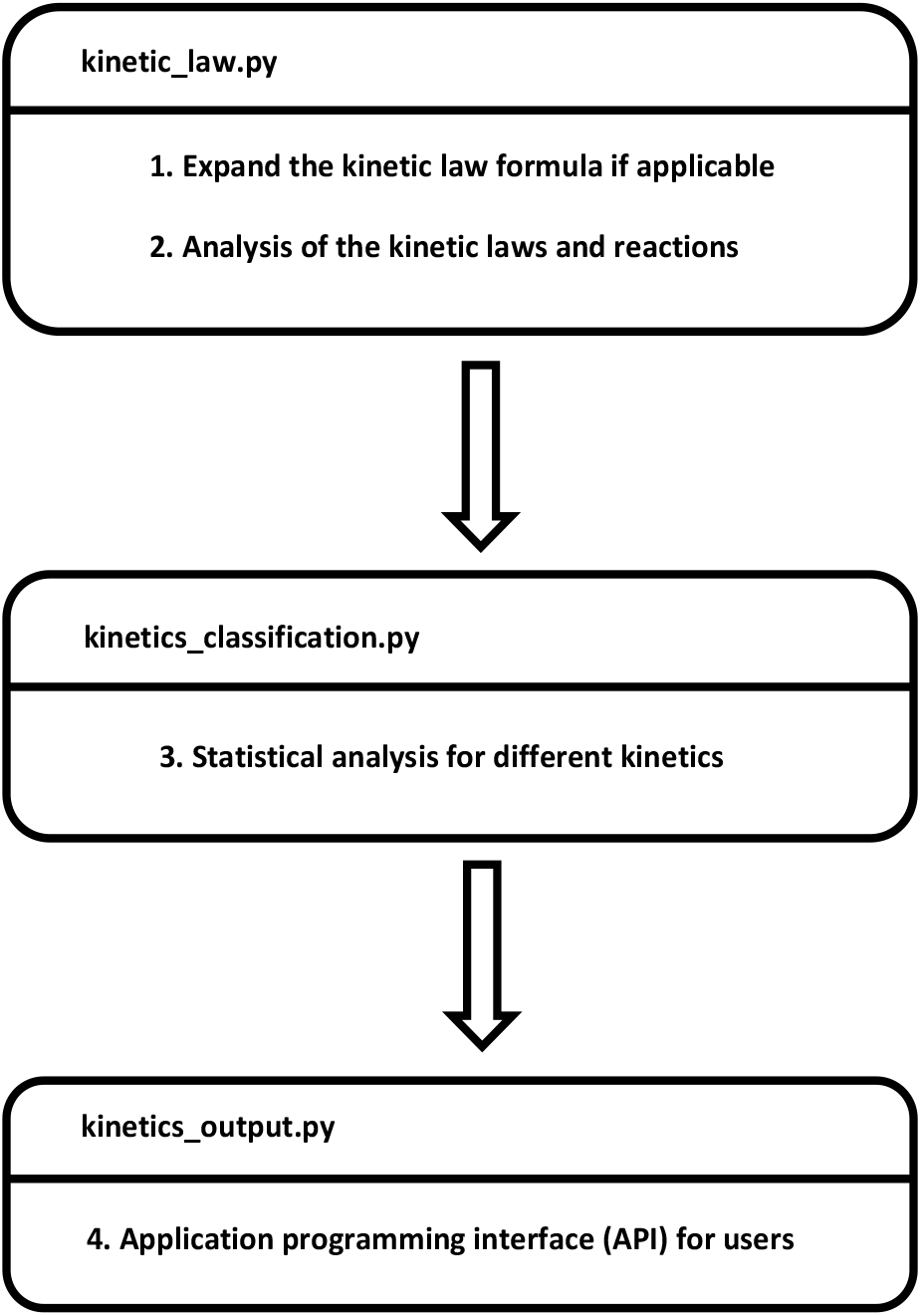
Code structure and workflow steps of SBMLKinetics.

#### 2.1 Expand the kinetic law formula if applicable

Before analyzing kinetics, I expanded the kinetic law formula if it was in the format with a function name. In the SBML files, some kinetic laws are expressed via specific function names. Therefore, the function names are supposed to get replaced by certain function bodies. For instance, the kinetic law is sometimes defined by *f* (*t*). Then, it is necessary to expand *f* (*t*) to its explicit equation *at* + *b* based on the definition of *f* (*x*) = *ax* + *b* in the SBML file.

#### 2.2 Analysis of the kinetic laws and reactions

K types were determined by a collection of eight kinetics properties (K properties) listed below that were ascertained from analyzing the reaction’s kinetic law.

a. the number of species in the kinetic law;
b. whether the kinetic law was a single product of terms;
c. whether the kinetic law was the difference between two products of terms;
d. whether the first (and the second) product of terms contained species as all the reactants (and products);
e. whether the kinetic law was in the format of Michaelis-Menten expressions with a single reactant in the numerator;
f. whether the kinetic law was in the format of Michaelis-Menten expressions with the product of a single reactant and another species in the numerator;
g. whether the kinetic law was in the format of Hill equation;
h. whether the kinetic law was in the fraction format with at least one species in the denominator.

The logic for determining the **K type** based on K properties was described in Table 3. For example, if the number of species in the kinetic law was zero according to K property **a**, then the **K type** was ZERO. If both K properties **b** and **d** held, then the **K type** was UNDR. The analysis proceeded column by column in Table 3. Note that the classifications were mutually exclusive since all pairs of columns differed in at least one non-empty row. For example, UNDR and UNMO differed in the K property **d**. To illustrate the above analysis, consider the kinetic law cell*·*CP*·*k9 of the reaction CP → C2 in the second row of Table 1. The K property **b** (it was a single product of terms) and **d** (all reactants were in the kinetic law) both held. Therefore, I saw that its **K type** was UNDR based on Table 3.

**Table 3.** The determination of kinetics types (K types) by kinetics properties (K properties).

| K properties | ZERO | UNDR | UNMO | BIDR | BIMO | MM | MMCAT | HILL | FR |
| --- | --- | --- | --- | --- | --- | --- | --- | --- | --- |
| a. | = 0 | > 0 | > 0 | > 0 | > 0 | 1 | 2 | 1 | > 0 |
| b. |  | Yes | Yes |  |  |  |  |  |  |
| c. |  |  |  | Yes | Yes |  |  |  |  |
| d. |  | Yes | No | Yes | No |  |  |  |  |
| e. |  |  |  |  |  | Yes |  |  | No |
| f. |  |  |  |  |  |  | Yes |  | No |
| g. |  |  |  |  |  |  |  | Yes | No |
| h. |  |  |  |  |  | Yes | Yes | Yes | Yes |

To classify the R types, I considered the number of distinct reactants and products. **R type** was quantitatively represented by the number of reactants (R = 0, 1, 2, >2) and products (P = 0, 1, 2, >2).

#### 2.3 Statistical analysis for different kinetics

After analyzing the kinetic laws with the strategies in steps 1 and 2, I did a statistical analysis to obtain the following information.

- Detailed Information for each reaction including BioModel identifier number, reaction id, reaction, kinetic law, and **K type**.
- Detailed information for each BioModel including BioModel identifier number, the number of reactions, and the distributions of ten K types per model.

#### 2.4 Application programming interface (API) for users

Finally, users could access the results via APIs in the following perspectives.

- Query distributions of **K type** and **R type**.
- Query elements of (top) **K type** and **R type**.
- Plots for the distributions of **K type** and **R type**.

All the API details were documented and available at GitHub (https://sunnyxu.github.io/SBMLKinetics/).

### 3. Uniqueness

The cross-platform Python-based tool, SBMLKinetics, constructed reaction classifications by inputting a collection of SBML models and outputting the probability of each **2DK** class (combinations of **K type** and **R type**). The kinetic analysis was based on python-libSBML [28], which supports SBML in levels 1-3 [29, 30, 31]. Therefore, SBMLKinetics could rely on any SBML database in levels 1-3.

To compare with SBMLsqueezer 2 [21] which is the only tool for large-scale biochemical kinetic models, I summarized its major differences from SBMLKinetics in Table 4. SBMLsqueezer 2 uses a classification scheme, called *de novo* creation method, that “considers the annotation of all participating reactants, products, and regulators” [24]. It contains a connection to the kinetics database to recommend kinetic laws. In detail, its extraction of kinetic laws from the SABIO-RK [26] database is based on the similarity of annotations which cannot always yield results [21]. Briefly, SBMLsqueezer uses a creative method to classify kinetics that relies heavily on the information from annotations. However, annotations are not always available in SBML files in model repositories such as BioModels.

**Table 4.** Comparison between SBMLKinetics and SBMLsqueezer 2.

| Major differences | SBMLKinetics | SBMLsqueezer 2 |
| --- | --- | --- |
| Interface | command line | various frameworks |
| Annotations | independent | dependent |
| Classification scheme | **2DK** | *de novo* creation |
| Recommending kinetic laws | fully *data-driven* | mainly *theory-driven* |
| Reliability of database | any SBML database in levels 1-3 [29, 30, 31] | SABIO-RK [26] |

The tool of SBMLKinetics made use of an annotation-independent classification scheme to deal with the kinetics analysis based on the SBML database, which could include all the current and future information from the SBML model database and experimental results. The categories in the *de novo* creation of SBMLsqueezer 2 are not exclusive to each other as the **2DK** of SBMLKinetics. In comparison with the *de novo* creation, **2DK** had covered all the categories of the *de novo* creation except the perspectives of reversible or irreversible reactions. Table 5 provided more details.

**Table 5.** Comparison between *de novo* creation in SBMLsqueezer 2 and **2DK** in SBMLKinetics.

| **de novo ceation** | **2DK** | specific descriptions if applicable |
| --- | --- | --- |
| Non-enzyme reactions | K type | ZERO, UNDR, UNMO, BIDR, BIMO |
| Uni-uni enzyme reactions | K type | MM, MMCAT, HILL |
| Bi-uni enzyme reactions | K type | FR |
| Bi-bi enzyme reactions | K type | FR |
| Arbitrary enzyme reactions | K type | FR |
| Modulated reactions | K type | UNMO, BIMO, MM, MMCAT, HILL |
| Gene-regulatory processes | R type or K type | R = 0 or Hill/FR |
| Integer stoichiometry reactions | R type |  |
| Zeroth reactant order reactions | R type | R = 0 |
| Zeroth product order reactions | R type | P = 0 |
| irreversible reactions | – |  |
| reversible reactions | – |  |

## Results and Validation

I considered a classification scheme to be useful if: (a) the classification categories provided useful insights, and (b) the scheme could classify a large fraction of items. The discussion in the Methods section supported (a) that **2DK** classifications aligned well with easily identifiable properties of reactions. Here, I addressed (b). In particular, I calculated the coverage of **2DK**, namely the fraction of the kinetic laws of reactions in curated BioModels that could be classified using **2DK**, which turned out to be approximately 95%. In the following subsections, I presented more detailed information about **K type** and **R type** distributions.

### 1. Kinetics type distribution

There was possible kinetics like zeroth order, mass action, Michaelis-Menten, Hill kinetics, and others. See the section Methods for details. Figure 2 characterized the kinetics used in the BioModels Database. The classification of the ten types was exclusive. Therefore, the sum of all the probabilities was one. Figure 2 showed the distribution of K types from 931 curated BioModels with 30,592 reactions. The top **K type** was UNDR, and 4.37% reactions were not classified. The relatively small number of unclassified reactions did suggest that the selected classification schemes were reasonable. The running time was 9.2 hours. All computations reported were done using an Intel i7 9700 processor running at 3.00 GHz with 32 GB RAM on Windows 10.

**Figure 2.**
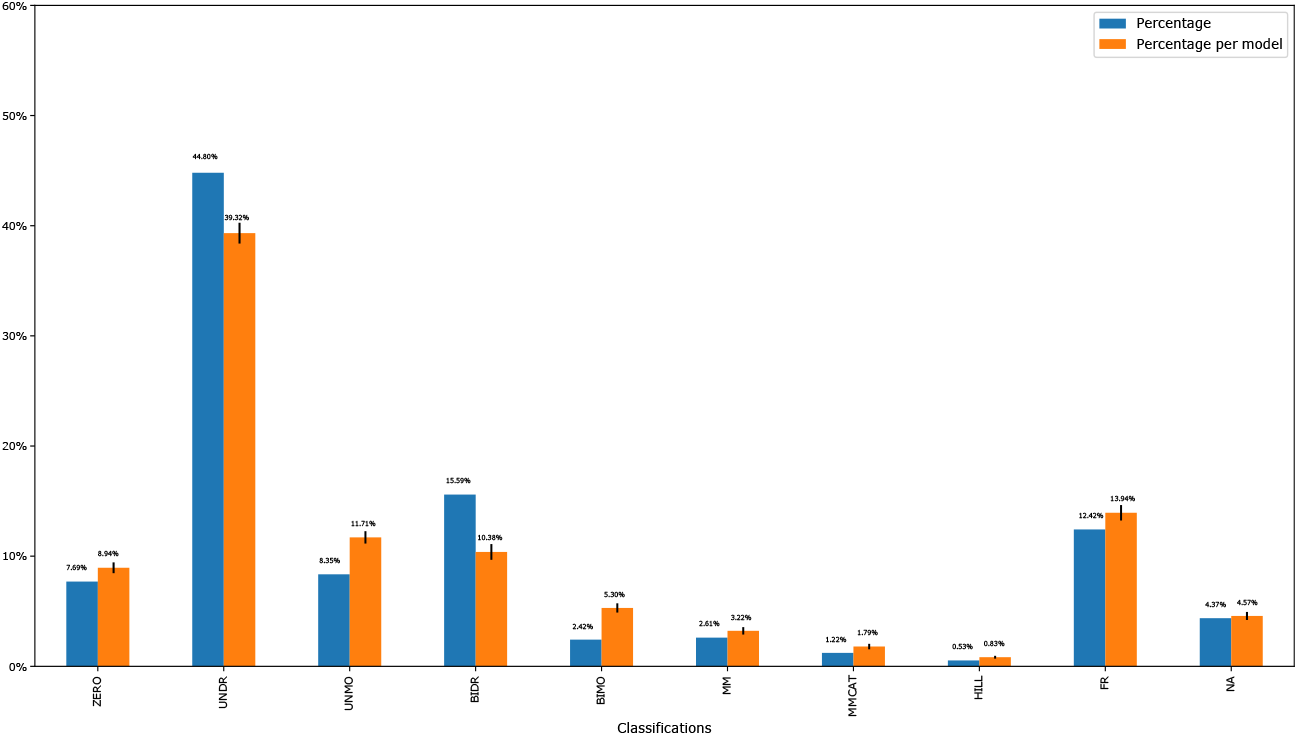
Kinetics type distribution. The blue bars investigated the kinetics type distribution of the average of all the reactions from all the models. The orange bars indicated the distribution of the average of reactions for each model. The trend for the two color bars was qualitatively similar. The error bars on the top of the orange bars represented the uncertainty among different models.

### 2. Reaction type distribution

To classify the reaction types (R types), I considered the number of distinct reactants and products. **R type** was quantitatively represented by the number of distinct reactants (R = 0, 1, 2, *>*2) and products (P = 0, 1, 2, *>*2). The classification of all the reaction types was exclusive. The sum of all the probabilities was one, which meant that I covered all the curated BioModels from the perspective of **R type**. Figure 3 showed the distribution of R types from 931 curated BioModels with 30,592 reactions. The top **R type** was with a single reactant and a single product.

**Figure 3.**
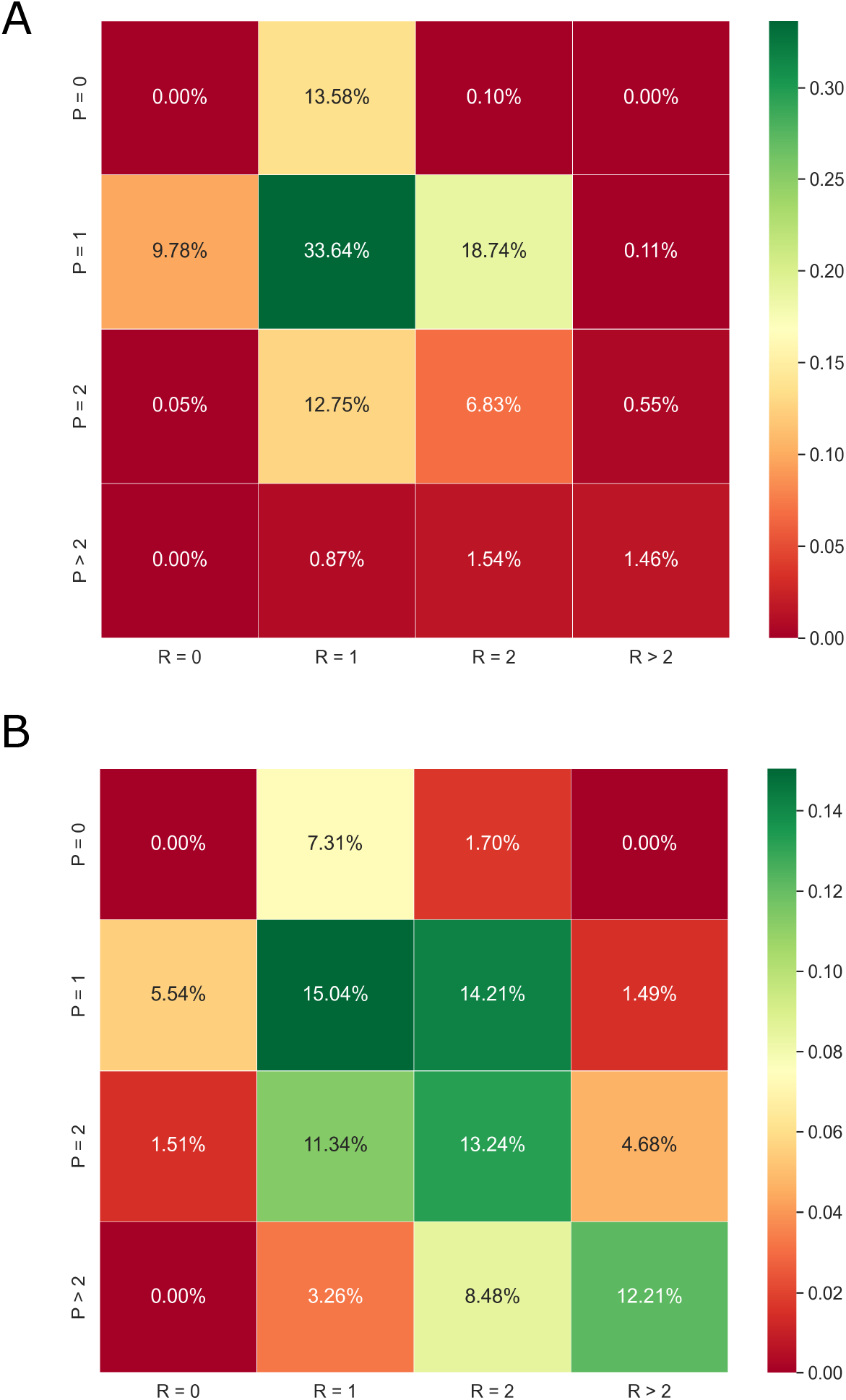
Reaction type distribution. (A) investigated the reaction type distribution of the average of all the reactions from all the models. (B) indicated the distribution of the average of reactions for each model. The trend for the two subplots was qualitatively similar.

There are several models in the Database, and there are several reactions in each model. In Figure 2, the blue bars investigated the **K type** distribution of the average of all the reactions from all the models. In detail, I counted the number of reactions in all the models as *R*_*all-models*_. Then, I counted the percentage of reactions for each **K type** among all the reactions (*R*_*all-models*_). The orange bars indicated the distribution of the average of reactions for each model. The error bars on the top of the orange bars represented the uncertainty among different models, which was relatively small. In detail, I first counted the number of reactions in each model as *R*_*per-model*_. Then, I counted the percentage of reactions for each **K type** among the reactions in the certain model (*R*_*per-model*_). The value of the distribution was the average of the percentage among all the models. The error bar was the standard error among all the models. For each **K type** distribution in Figure 2, the trend for the two color bars was qualitatively similar with relatively small error bars. Therefore, the statistics were meaningful for the **K type** distributions. In Figure 3, the trend for the two subplots was qualitatively similar which did not depend on a certain model. Therefore, the **R type** distribution was meaningful too.

## Applications and Discussion

**2DK** could provide a *data-driven* approach to recommending kinetic laws and providing an alert of unusual kinetic laws. I applied the **2DK** method and the tool SBMLKinetics to the BioModels Database as an example of analyzing kinetics from different reaction networks, although the tool could be applied to any repository of SBML kinetics models. The examples could illustrate the potential applications of the method and how to use the tool. In the following subsections, I indicated how the variety of **K type** distributions depended on different R types. In addition, I compared two subsets of BioModels including signaling networks and metabolic networks.

### 1. Kinetics type distributions for different reaction types

I investigated the distribution of kinetics based on reaction types and found substantial differences among them (Figure 4). Focusing on the four types in the center with the most reactions involved (R = 1, P = 1; R = 2, P = 1; R = 1, P = 2; R = 2, P = 2), these four subplots were quite different from each other. The top **K type** with one reactant was UNDR significantly, while the distributions became more evenly with two reactants.

**Figure 4.**
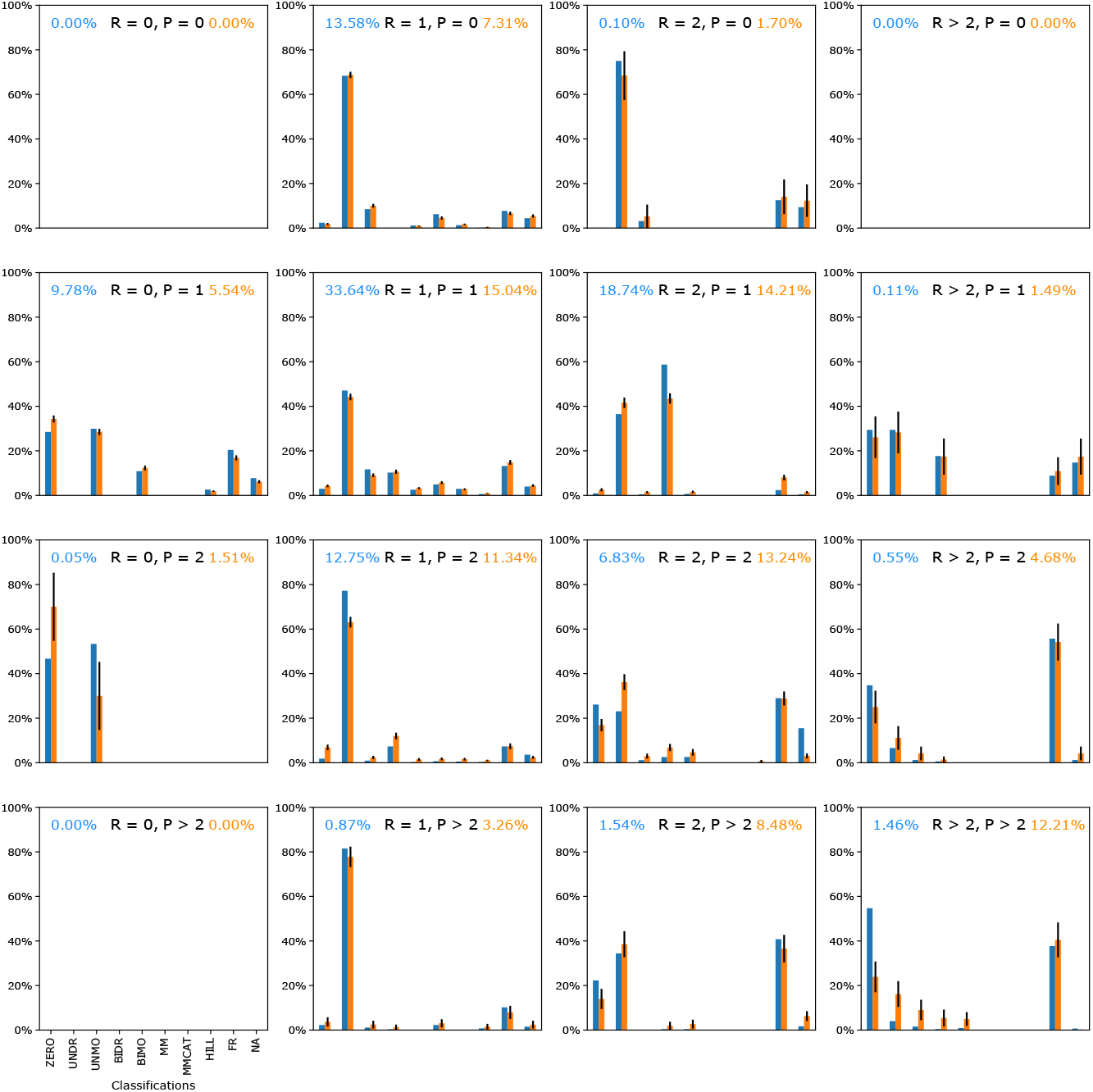
Kinetics type distribution for different reaction types. For each reaction type, represented by the number of distinct reactants and products (marked as the black text in each subplot), a kinetics type distribution was given. The blue text in the top left corner represented the percentage of reactions involved in a certain reaction type. The orange text in the top right corner represented the percentage of reactions per model. The blue bars investigated the kinetics type distribution of the average of all the reactions from all the models. The orange bars indicated the distribution of the average of reactions for each model. The trend for color bars was qualitatively similar for each reaction type. The error bars on the top of the orange bars represented the uncertainty among different models.

Figure 4 investigated the significant differences in **K type** distributions among R types. The blue text in the top left corner represented the percentage of reactions involved in the certain **R type**, which was more clearly illustrated in Figure 3A. The orange text in the top right corner represented the percentage of reactions per model, which was more clearly indicated in Figure 3B. Following the observation of the blue and orange texts, the most frequent six R types were (R = 1, P = 0; R = 0, P = 1; R = 1, P = 1; R = 2, P = 1; R = 1, P = 2; R = 2, P = 2). In Figure 4, the **K type** distributions were non-symmetric depending on different R types, which meat the effects from the number of distinct reactants and products were not equal. Based on Figure 4, researchers could tell which **R type** had the highest or lowest **K type**. For example, the **R type** with R = 2, P = 1 had the highest BIDR among all the R types. While there was the highest NA from the **R type** with R *>* 2, P = 1. It was also interesting to compare per column (with the same number of reactants) or row (with the same number of products). For instance, the third row indicated that the FR became more frequent as the number of reactants increased with a constant number of the product (P = 2).

### 2. Comparison between different subsets of BioModels

This characterization considered the biology being studied as well as the nature of the reactions themselves. For instance, I compared the **K type** distributions for signaling and metabolic networks in the BioModels Database as an example. For the details of selecting the two subsets, see the Availability of data and materials under the Declarations section. I found substantial differences between the two types of networks in Figure 5. Figure 5A showed the distribution of K types from 288 signaling networks with 13,572 reactions. The top **K type** was UNDR, and 1.84% reactions were not classified. The running time was 6.9 hours. Figure 5B showed the distribution of K types from 168 metabolic networks with 6,904 reactions. The top **K type** was FR, and 9.85% reactions were not classified. The running time was 8.1 hours. The **K type** distribution of signaling networks was similar to the whole curated BioModels in Figure 2. However, the metabolic networks were quite different with high frequencies of ZERO and FR.

**Figure 5.**
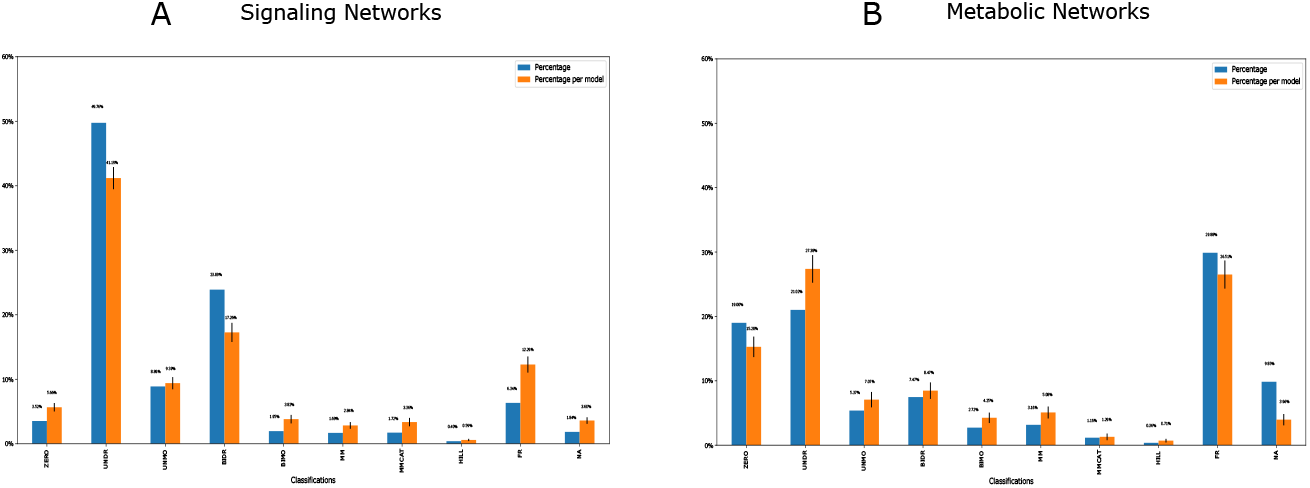
Kinetics type distribution for signaling networks (A) and metabolic networks (B). The blue bars investigated the kinetics type distribution of the average of all the reactions from all the models. The orange bars indicated the distribution of the average of reactions for each model. The error bars on the top of the orange bars represented the uncertainty among different models. The trend for the two color bars was qualitatively similar for each subplot. However, there was a significant difference in the kinetics type distributions between the signaling and metabolic networks.

The significant difference between signaling networks and metabolic networks could come from regulatory interactions. The high frequency of ZERO and FR from metabolic networks was the kinetics which could be very simple without any species inside the kinetic law or a very complicated kinetic law that had a fractional representation.

The different **K type** distributions among different data sets, if applied to synthetic random reaction networks, could provide more representative kinetic laws for these studies. As an example to indicate how this method could contribute to generating synthetic reaction networks, I made use of the results from the BioModels Database as an example. However, users could generate a distribution based on their database for further research. Based on the result of Figure 2 from curated BioModels, users could select UNDR with the highest probability as the general mass action kinetics to model synthetic reaction networks. Users could also follow its distribution to assign the probability for each type of kinetics while generating random synthetic reaction networks. In detail, users could generate reaction networks by assigning the probability of certain kinetics, i.e., ZERO, UNDR, UNMO, etc. Researchers commonly use simplified mass action kinetics as a first trial while modeling synthetic reaction networks [1], therefore, users could choose a certain distribution referring to Figure 4 if they had already known the **R type** information. For example, if there were one reactant and one product in the reaction, UNDR was the most common kinetics to choose. While BIDR was more likely to get chosen for the case with two reactants and one product. If users wanted to model synthetic signaling networks instead of metabolic networks, they could refer to the distributions of Figure 5A instead of Figure 5B.

## Conclusions

SBMLKinetics could analyze any data sets with SBML files as input and generate the analysis of kinetics. Here, I used the BioModels Database as an SBML database example to show the validation and potential applications of **2DK**. However, users could use their experimental data or another model database to look for natural biological properties. See the Availability of data and materials under the Declarations section.

Therefore, **2DK** could provide a *data-driven* approach to recommending kinetic laws and providing an alert of unusual kinetic laws. This work had the potential application to computational studies based on random reaction networks so that more representative kinetic laws were used in these studies. As I had discussed the relationship between **K type** and **R type**, this tool could suggest the probabilities of **K type** for each **R type**. I could also observe how the **K type** distributions vary according to different R types. The methods could make suggestions to choose **K type** distributions depending on different models, for instance, signaling vs. metabolic networks. Statistically, the probability distributions of K types could be a prior for kinetic law generations, as Bayesian inference has been discussed about kinetics before [32, 33].

What if the kinetic laws inside a model were significantly different from the distributions of the **K type** or **R type** calculated by the reference models? It could mean that the model had errors in its kinetics. At a minimum, the presence of such differences should be explored by the modeler. But such differences might also reflect substantially different biology or chemistry that was under study.

SBMLKinetics could be useful to pursue biological insights from updating models or experimental databases. For example, it is essential to compare the kinetics among different data sets, i.e., different organisms. The future potential work could be further applications of **2DK**. Applications could include a complete recommending system for specific kinetic laws, detecting errors in kinetic laws, and seeking the biology insights of models based on such classifications.

## Declarations

## Acknowledgements

J.X. thanks Joseph L. Hellerstein for introducing the problems considered in this paper, many stimulating discussions, his comments on the manuscript, and a grammar check. J.X. appreciates Herbert M. Sauro’s discussions regarding kinetics and synthetic reaction networks cited as the references, and for his comments on the early stage of the manuscript. J.X. also thanks Lucian Smith for the discussion about the kinetic law formula expansion based on libSBML. Finally, J.X. appreciates the valuable comments from the reviewers.

## Abbreviations

I. Kinetics type (**K type**)

**K type** includes ten types:

ZERO: Zeroth order
UNDR: Uni-directional mass action
UNMO: Uni-term with the moderator
BIDR: Bi-directional mass action
BIMO: Bi-terms with the moderator
MM: Michaelis-Menten kinetics without an explicit enzyme
MMCAT: Michaelis-Menten kinetics with an explicit enzyme
HILL: Hill kinetics
FR: Kinetic law in the fraction format other than MM, MMCAT, or HILL
NA: not classified

II. Reaction type (**R type**)

**R type** was quantitatively represented by the number of distinct reactants (R = 0, 1, 2, *>*2) and products (P = 0, 1, 2, *>*2).

## Availability of data and materials

SBMLKinetics was a publicly available Python package (https://pypi.org/project/SBMLKinetics/) and was licensed under the liberal MIT open-source license. The package was available for all major platforms. The source code had been deposited on GitHub (https://github.com/SunnyXu/SBMLKinetics). Users could install the package using the standard pip installation mechanism: pip install SBMLKinetics. The package was fully documented as https://sunnyxu.github.io/SBMLKinetics/. The specific version of the tool SBMLKinetics used for the analysis and results presented in the article was version 1.0.1.

As BioModels Database is changing and improving over time, all the data sets to produce the results and analysis in the article were available in the GitHub repository (https://github.com/SunnyXu/SBMLKinetics/tree/master/data). Initially, the metabolic datasets were downloaded from the BioModels Database website (https://www.ebi.ac.uk/biomodels/), by clicking on the “Manually Curated” button, selecting the “SBML” model format, and typing “AND “Metabolic” “ in the searching engine. For signaling networks, both “Signaling” and “Signalling” were considered.

## Ethics approval and consent to participate

The data that I used was obtained from the public datasets BioModels Database (https://www.ebi.ac.uk/biomodels/). Therefore, the ethics approval did not apply to this study.

## Consent for publication

Not applicable.

## Authors’ contributions

J.X. implemented the project, did the data analysis, and was the main developer of the Python package. J.X. wrote the manuscript.

